# Dual Role of Neuroplastin in Pancreatic β Cells: Regulating Insulin Secretion and Promoting Islet Inflammation

**DOI:** 10.1101/2023.09.08.556759

**Authors:** Rie Asada Kitamura, Devynn Hummel, Fumihiko Urano

## Abstract

Mesencephalic astrocyte-derived neurotrophic factor (MANF) is an endoplasmic reticulum (ER)-resident secretory protein that reduces inflammation and promotes proliferation in pancreatic β cells. Numerous studies have highlighted the potential of MANF as a therapeutic agent for diabetes mellitus (DM), making it essential to understand the mechanisms underlying MANF’s functions. In our previous search for a molecule that mediates MANF signaling, we identified Neuroplastin (NPTN) as a binding partner of MANF that localizes on the cell surface. However, the roles of NPTN in pancreatic β cells remain unclear. In this study, we generated β cell-specific *Nptn* knockout (KO) mice and conducted metabolic characterization. NPTN deficiency improved glucose tolerance by increasing insulin secretion and β cell mass in the pancreas. Moreover, proliferation and mitochondrial numbers in β cells increased in *Nptn* KO islets. These phenotypes resulted from elevated cytosolic Ca^2+^ levels and subsequent activation of downstream molecules. Simultaneously, we demonstrated that NPTN induces the expression of proinflammatory cytokines via the TRAF6-NF-κB axis in β cells. Additionally, NPTN deficiency conferred resistance to STZ-induced diabetic phenotypes. Finally, exogenous MANF treatment in islets or β cells led to similar phenotypes as those observed in NPTN-deficient models. These results indicate that NPTN plays important roles in the regulation of insulin secretion, proliferation, and mitochondrial quantity, as well as pro-inflammatory responses, which are antagonized by MANF treatment. Thus, targeting the MANF-NPTN interaction may lead to a novel treatment for improving β cell functions in diabetes mellitus.

**Significance statement:** Mesencephalic astrocyte-derived neurotrophic factor (MANF) is an endoplasmic reticulum (ER)-resident small secretory protein that has the potential as therapeutic agent for various diseases related to inflammation and ER stress, such as Type 1 diabetes mellitus. Our work shed light on the roles of a binding partner protein of MANF, Neuroplastin (NPTN), in pancreatic β cells. We demonstrated NPTN regulates Ca^2+^ dynamics as well as inflammation in pancreatic β cells. NPTN deficiency caused improved insulin secretion as well as the resistance to Type 1 diabetic phenotypes. We also found out that MANF treatment leads to similar phenotypes observed in NPTN deficient models through antagonizing NPTN’s functions. Overall, our results provide a new insight into treatment for improving β cell functions in diabetes mellitus.

## Introduction

Pancreatic β cells are uniquely responsible for insulin production and secretion to counteract elevated blood glucose levels and are susceptible to metabolic and cytotoxic assaults. β cell dysfunction and death ultimately lead to diabetes mellitus (DM), a set of metabolic disorders characterized by dysregulated glucose homeostasis. In the etiology of type 1 diabetes mellitus (T1D), β cells are selectively targeted by autoimmunity, resulting in islet inflammation and reduced insulin production and secretion. Additionally, endoplasmic reticulum (ER) stress occurs in β cells during T1D pathogenesis, further accelerating dysfunction and apoptosis of these cells. (1, 2).

Mesencephalic astrocyte-derived neurotrophic factor (MANF) is an endoplasmic reticulum (ER)-resident small secretory protein that is emerging as a beneficial molecule against the pathophysiology of various diseases related to inflammation and ER stress. This includes genetic disorders such as Wolfram syndrome (3-5). Under normal conditions, MANF is retained within the ER by binding to the ER-resident molecular chaperone, BiP (6, 7). Inflammatory stimuli and ER stress cause the release of MANF from BiP, resulting in the secretion of a certain amount of MANF proteins into extracellular spaces (7-9). Simultaneously, MANF expression is also induced by unfolded protein response (UPR) signals (10-12). Accumulating evidence suggests that both ER-resident and extracellular MANF mitigate inflammation and ER stress in a variety of tissues, thereby protecting these tissues from the development and progression of disease conditions (3, 4). Notably, whole-body and pancreatic β cell-specific *Manf* KO mice exhibited increased ER stress and β cell death, leading to the spontaneous development of diabetes (13, 14). On the other hand, local overexpression of *Manf* in pancreatic β cells inhibits or reverses diabetic phenotypes in either streptozotocin (STZ)-injected rodent models or genetic disorder models (13). In addition to its cytoprotective effects, MANF promotes the proliferation of β cells (5, 12-14). Moreover, it has been reported that the level of circulating MANF in the blood is elevated in patients with both type 1 diabetes (T1D) and type 2 diabetes (T2D) (15, 16). These experimental and clinical considerations suggest that MANF is a highly attractive molecule with potential for therapeutic applications.

In the context of the molecular mechanism by which MANF exerts its cytoprotective effects, some studies have demonstrated that ER-resident MANF prevents the oligomerization of ER stress transducers, such as IRE1, which would otherwise provoke an apoptotic signal (17, 18). Another study suggested that MANF contributes to protein folding by functioning as a nucleotide exchange inhibitor for BiP (18). As for secreted MANF, a study focusing on the N-terminal saposin-like domain of MANF, which has a high affinity for lipid sulfatide present in outer-cell membrane leaflets, indicated that MANF is taken up through endocytosis and subsequently transported into the ER lumen (19). Additionally, it has been shown that treatment with recombinant MANF proteins inhibits the activation of NF-κB, a major transcription factor of inflammatory genes (12). However, the precise mechanism, specifically how extracellular MANF modulates NF-κB signaling within cells, remains unclear.

In pursuit of a molecule that mediates the extracellular signal of MANF into the cellular signal, we previously identified a binding partner of MANF on the cell surface, Neuroplastin (NPTN), using a proteomics strategy (20). Our previous in vitro studies demonstrated that NPTN induces the expression of pro-inflammatory cytokine genes by activating NF-κB signaling, which is antagonized by a direct binding of MANF to NPTN (20). NPTN is an immunoglobulin superfamily glycoprotein, with two isoforms, Np65 and Np55 (21, 22). Np65 is specifically expressed in neurons of the brain, particularly the forebrain, and plays important roles in neuronal functions (23-25). On the other hand, Np55 is more ubiquitously expressed compared to Np65 (23-25). Given that MANF is indispensable for pancreatic β cell survival, NPTN is likely to be an important molecule not only in neurons but also in other types of cells in peripheral tissues, especially pancreatic β cells. However, the role of NPTN in pancreatic β cells remains unclear.

Here, we generate pancreatic β cell-specific *Nptn* KO mice and *Nptn* KO Rat insulinoma cell line INS-1 832/13 to investigate the roles of NPTN in pancreatic β cells. We identify NPTN as an essential molecule for Ca^2+^ homeostasis, which leads to proper insulin secretion and proliferation of β cells. Furthermore, we demonstrate that NPTN contributes to the development of STZ-induced diabetic phenotypes, and that MANF antagonizes these NPTN roles.

## Results

### NPTN deficiency improves glucose tolerance by increasing glucose-stimulated insulin secretion, proliferation, and mitochondrial quantity in β cells

To investigate the role of NPTN in pancreatic β cells, we generated pancreatic β cell-specific *Nptn* knockout mice by breeding *Nptn^flox/flox^* mice with *Ins1^Cre^* mice (*Nptn*^f/f^; *Ins1^Cre^*) (26) (Figure S1A). We confirmed the robust reduction of NPTN expression in islets isolated form *Nptn*^f/f^; *Ins1^Cre^* mice (Figure S1B). Immunofluorescence revealed that NPTN is expressed in insulin-positive β cells as well as glucagon-positive α cells (Figure 1A) and only the NPTN signals of β cells were depleted in *Nptn*^f/f^; *Ins1^Cre^*pancreas sections (Figure 1A). We first identified metabolic phenotypes in *Nptn*^f/f^; *Ins1^Cre^* mice. An intraperitoneal glucose tolerance test (IP-GTT) demonstrated that *Nptn*^f/f^; *Ins1^Cre^* mice improved glucose tolerance in both male and females at 7 weeks old (Figure 1B and Figure S2A). On the other hand, an intraperitoneal insulin tolerance test (IP-ITT) curves were similar regardless of genotype (Figure 1C and Figure S2B). Elevation of serum insulin levels following glucose injection were higher in *Nptn*^f/f^; *Ins1^Cre^* mice as compared to *Nptn*^f/f^ mice (Figure 1D and Figure S2C). Basal serum insulin level was also increased in the male *Nptn*^f/f^; *Ins1^Cre^* (Figure 1D). Static glucose stimulated insulin secretion (GSIS) assay showed that the functional capacity of islets isolated from *Nptn*^f/f^; *Ins1^Cre^* mice was improved as compared to *Nptn*^f/f^ islets (Figure 1E), whereas insulin content was similar (Figure 2F). However, insulin content in whole pancreas was increased in both male and female *Nptn*^f/f^; *Ins1^Cre^* mice (Figure 2G and Figure S2D). Additionally, insulin-positive β cell mass was increased in *Nptn*^f/f^; *Ins1^Cre^* mice (Figure 1H and Figure S2E). These results suggested that NPTN deficiency increases GSIS as well as β cell mass of whole pancreas, which contributes to improved glucose tolerance in *Nptn*^f/f^; *Ins1^Cre^* mice.

**Figure 1.**
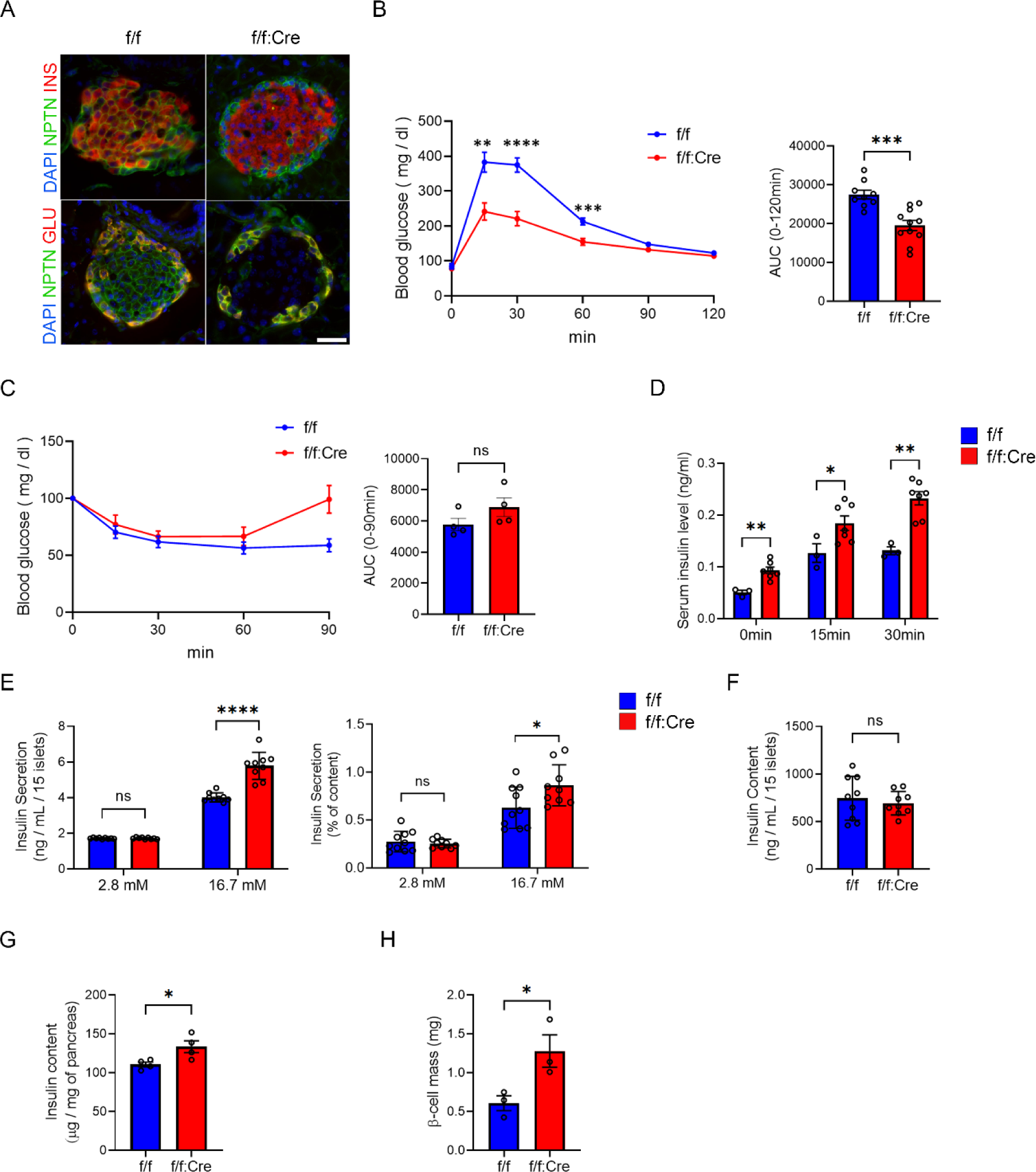
**NPTN deficiency increases glucose-stimulated insulin secretion (GSIS) and β cell mass** (A) Representative immunofluorescence images of NPTN, insulin, and glucagon in islets from *Nptn*^f/f^ or *Nptn*^f/f^; *Ins1^Cre^* mice. Scale bar: 25 μm. (B) (Left) IP-GTT with *Nptn*^f/f^ or *Nptn*^f/f^; *Ins1^Cre^* male mice. (Right) AUCs of the IP-GTT (*Nptn*^f/f^: n=9, *Nptn*^f/f^; *Ins1^Cre^*: n=11, **p<0.01, ***p<0.001 and ****p<0.0001 by unpaired *t*-test). (C) (Left) IP-ITT with *Nptn*^f/f^ or *Nptn*^f/f^; *Ins1^Cre^* male mice. (Right) AUCs of the IP-ITT (*Nptn*^f/f^: n=4, *Nptn*^f/f^; *Ins1^Cre^*: n=4). (D) Serum insulin levels following glucose injection in *Nptn*^f/f^ or *Nptn*^f/f^; *Ins1^Cre^*male mice (*Nptn*^f/f^: n=3, *Nptn*^f/f^; *Ins1^Cre^*: n=7, *p<0.05, **p<0.01 by unpaired *t*-test). (E) Static GSIS per islets (left) and normalized to insulin content (right) in primary islets form *Nptn*^f/f^ or *Nptn*^f/f^; *Ins1^Cre^*mice (*Nptn*^f/f^: n=10, *Nptn*^f/f^; *Ins1^Cre^*: n=9, *p<0.05 and ****p<0.0001 by unpaired *t*-test). (F) Insulin content in the islets used in (E). (G) Insulin content of whole pancreas from *Nptn*^f/f^ or *Nptn*^f/f^; *Ins1^Cre^* male mice (*Nptn*^f/f^: n=4, *Nptn*^f/f^; *Ins1^Cre^*: n=4, *p<0.05 by unpaired *t*-test). (H) β cell mass of pancreas in *Nptn*^f/f^ or *Nptn*^f/f^; *Ins1^Cre^* male mice (*Nptn*^f/f^: n=3, *Nptn*^f/f^; *Ins1^Cre^*: n=3, *p<0.05 by unpaired *t*-test). f/f: *Nptn*^f/f^, f/f:Cre: *Nptn*^f/f^; *Ins1^Cre^*, ns: not statistically significant.

**Figure 2.**
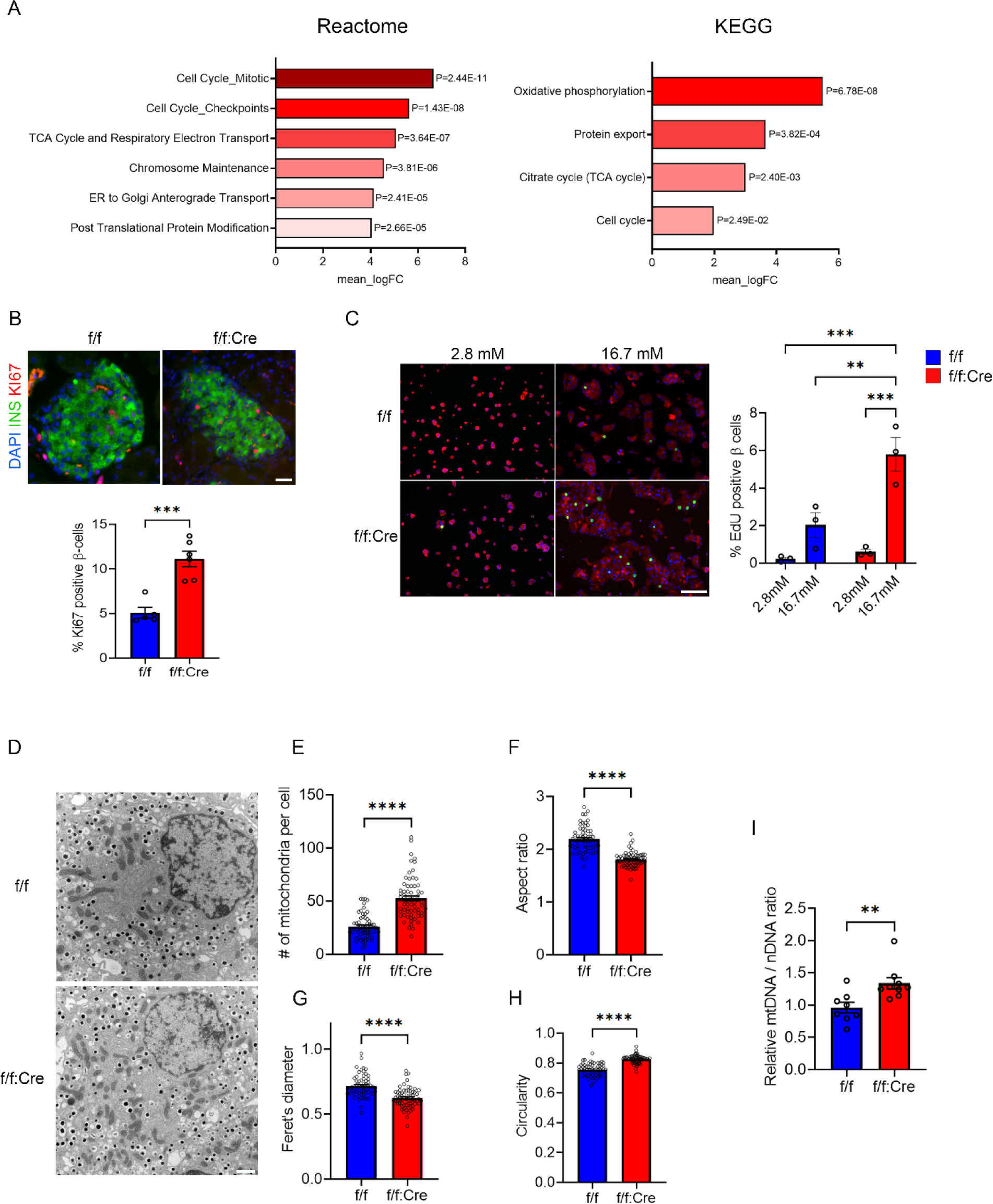
**NPTN deficiency increases proliferation and mitochondrial numbers in βcells** (A) Reactome and KEGG enrichment analyses for pathways upregulated in *Nptn*^f/f^; *Ins1^Cre^* islets (*Nptn*^f/f^: n=4, *Nptn*^f/f^; *Ins1^Cre^*: n=4). (B) (Upper) Representative immunofluorescence images of insulin and Ki67 in pancreas from *Nptn*^f/f^ or *Nptn*^f/f^; *Ins1^Cre^* mice. (Lower) A quantification of percentages of Ki67-positive β cells (*Nptn*^f/f^: n=5 (76 islets), *Nptn*^f/f^; *Ins1^Cre^*: n=6 (74 islets), ***p<0.001 by unpaired *t*-test). (C) (Left) Representative immunofluorescence images of EdU and insulin in *Nptn*^f/f^ or *Nptn*^f/f^; *Ins1^Cre^* islet cells cultured in the medium containing 2.8 mM or 16.7 mM glucose for 3 days. (Lower) A quantification of percentages of EdU-positive β cells (*Nptn*^f/f^, 2.8 mM: n=3 (7894 cells), *Nptn*^f/f^, 16.7 mM: n=3 (18260 cells), *Nptn*^f/f^; *Ins1^Cre^*, 2.8 mM: n=3 (6166 cells), *Nptn*^f/f^; *Ins1^Cre^*, 16.7 mM: n=3 (35698 cells), ***p<0.001 by two-way ANOVA). (D) Representative EM images of β cells in in *Nptn*^f/f^ or *Nptn*^f/f^; *Ins1^Cre^* islets. (E-H) A quantification of (E) mitochondrial number per cell (F) aspect ratio, (G) Feret’s diameter, and (H) circularity (*Nptn*^f/f^, n=61 cells, *Nptn*^f/f^; *Ins1^Cre^*, n=68 cells from 3 mice for each genotype. ****p<0.0001 by unpaired *t*-test). (I) Relative mitochondrial DNA (mtDNA) copy numbers normalized to nuclear DNA (nDNA) measured by qPCR analysis in islets from *Nptn*^f/f^ or *Nptn*^f/f^; *Ins1^Cre^*mice (*Nptn*^f/f^, n=8 cells, *Nptn*^f/f^; *Ins1^Cre^*, n=9, **p<0.01 by unpaired *t*-test). f/f: *Nptn*^f/f^, f/f:Cre: *Nptn*^f/f^; *Ins1^Cre^*, ns: not statistically significant.

To better understand the impact of *Nptn* knockout in β cells, we next performed bulk RNA-sequencing (RNA-seq) with islets isolated from *Nptn*^f/f^ and *Nptn*^f/f^; *Ins1^Cre^* mice at 7 weeks old. Enrichment analyses revealed the gene sets of cell cycle were significantly upregulated in *Nptn*^f/f^; *Ins1^Cre^* islets (Figure 2A). Interestingly, the gene sets pertaining metabolisms in mitochondria was also upregulated in *Nptn*^f/f^; *Ins1^Cre^* islets (Figure 2A and Figure S3). In addition, weighted gene correlation network analysis (WGCNA) (27) showed higher expression of gene set related to citrate cycle (TCA cycle) in *Nptn*^f/f^; *Ins1^Cre^* islets (Figure S3). These findings led us to hypothesize that NPTN deficiency increases proliferation and mitochondrial function or numbers in β cells. As predicted, the percentage of the insulin-positive cells expressing a proliferation marker, Ki67, was higher in *Nptn*^f/f^; *Ins1^Cre^* pancreatic sections (Figure 2B). It is known that the proliferation of rodent and human β cells is induced by high glucose stimulation (28-33). Consistent with the previous studies, our EdU cell proliferation assay using dispersed islet cells revealed that β cell proliferation was induced by high glucose stimulation (Figure 2C). However, the magnitude of induction was greater in *Nptn*^f/f^; *Ins1^Cre^* β cells (Figure 2C), indicating that NPTN deficiency leads to increased β cell proliferation. In addition, transmission electron microscopic (TEM) analyses showed that mitochondria were of greater abundance in *Nptn*^f/f^; *Ins1^Cre^* β cells as compared to *Nptn*^f/f^ β cells (Figure 2D and E). Interestingly, the mitochondria in *Nptn*^f/f^; *Ins1^Cre^* β cells displayed significantly shorter and more circular morphologies, as indicated by lower aspect ratios (Figure 2D and F), shorter feret’s diameters (Figure 2D and G), and circularity values being closer to 1 (Figure 2D and H), compared to that in *Nptn*^f/f^ β cells. These results imply that mitochondrial biogenesis or fission could be increased in *Nptn*^f/f^; *Ins1^Cre^* β cells. Similarly, mitochondrial DNA copy number, which reflects an abundance of mitochondria and β cell functional capacity (34-36), was increased in *Nptn*^f/f^; *Ins1^Cre^* islets (Figure 2I). On the other hand, the expression of β cell key genes in islets were similar regardless genotypes (Figure S4). In summary, NPTN deficiency in β cells causes an increase of cell proliferation and mitochondria number, leading to the improved metabolic output.

### NPTN is required for cytosolic Ca^2+^ homeostasis by regulating PMCA2 protein levels at the cell surface

To investigate the factor causing the phenotypes identified in *Nptn*^f/f^; *Ins1^Cre^* mice, we focused on cytosolic Ca^2+^ level. In pancreatic β cells, it is well known that an elevation of cytosolic Ca^2+^ level in response to glucose stimulation is crucial for insulin secretion as well as β cell proliferation (28, 31-33, 37-40). In addition, several studies demonstrated that mitochondria biogenesis and dynamics are induced by molecules functioning downstream of elevated cytosolic Ca^2+^ level (41-43). We, therefore, measured cytosolic Ca^2+^ levels in *Nptn*^f/f^ and *Nptn*^f/f^; *Ins1^Cre^* islet cells. *Nptn*^f/f^; *Ins1^Cre^* cells showed higher basal cytosolic Ca^2+^ levels as compared to *Nptn*^f/f^ cells (Figure 3A). Fold change of glucose-stimulated elevation of cytosolic Ca^2+^ levels was also higher in *Nptn*^f/f^; *Ins1^Cre^* cells (Figure 3B). Interestingly, NPTN has been reported to stabilize plasma membrane ATPases (PMCAs) through protein–protein interaction, which plays an important role for regulating cytosolic Ca^2+^ homeostasis (44-46). PMCAs are comprised of four isoforms (PMCA1-4), with PMCA2 being the most abundant isoform at the protein levels in pancreatic β cells (47, 48). As expected, PMCA2 protein level was significantly reduced in *Nptn*^f/f^; *Ins1^Cre^* islets (Figure 3C), whereas *Pmca2* mRNA level was similar regardless of genotypes (Figure S5A). We also confirmed the significant reduction of PMCA2 protein levels in *Nptn* KO INS1 832/13 cells (Figure S5B). Similar with the previous studies (44-46), the interaction between NPTN and PMCA2 was demonstrated in INS1 832/13 cells by immunoprecipitation (Figure 3D). Nuclear factor of activated T cells 2 (NFATc2) is one of the key transcription factors regulating β cell proliferation (49-51). The nuclear translocation of NFATc2 is regulated by calcineurin-mediated dephosphorylation, which is activated by elevated cytosolic Ca^2+^ levels (52, 53). Consistent with our EdU cell proliferation assay, the expression of NFATc2 target gene related to β cell proliferation (50), *Ccne2*, was significantly upregulated in response to high glucose stimulation in both *Nptn*^f/f^ and *Nptn*^f/f^; *Ins1^Cre^* islets (Figure 3E). Interestingly, the expressions of other proliferation-related NFATc2 target genes, *Ccna2* and *Cdk2*, were also significantly increased in glucose-stimulated *Nptn*^f/f^; *Ins1^Cre^* islets (Figure 3E). Moreover, the expression of *Pcna* was higher in *Nptn*^f/f^; *Ins1^Cre^* islets compared to *Nptn*^f/f^ islets (Figure 3E), suggesting that genes involved in β cell proliferation are more transcriptionally induced by NFATc2 in *Nptn*^f/f^; *Ins1^Cre^* islets. Calcineurin also dephosphorylates Ser 637 of DRP1, which is an important event for an increase in the mitochondrial fission rate (42). Consistent with our TEM analysis, phosphorylation levels of DRP1 Ser 637 were reduced in *Nptn*^f/f^; *Ins1^Cre^* islets (Figure 3F). Taken together, NPTN deficiency leads to higher basal and more dynamic cytosolic Ca^2+^ levels due to reduced PMCA2 proteins at expression at plasma membrane. Increased cytosolic Ca^2+^ promote calcineurin-mediated dephosphorylation of NFATc2 and DRP1 Ser 637 contributing to increases in β cell proliferation and mitochondrial fission, respectively.

**Figure 3.**
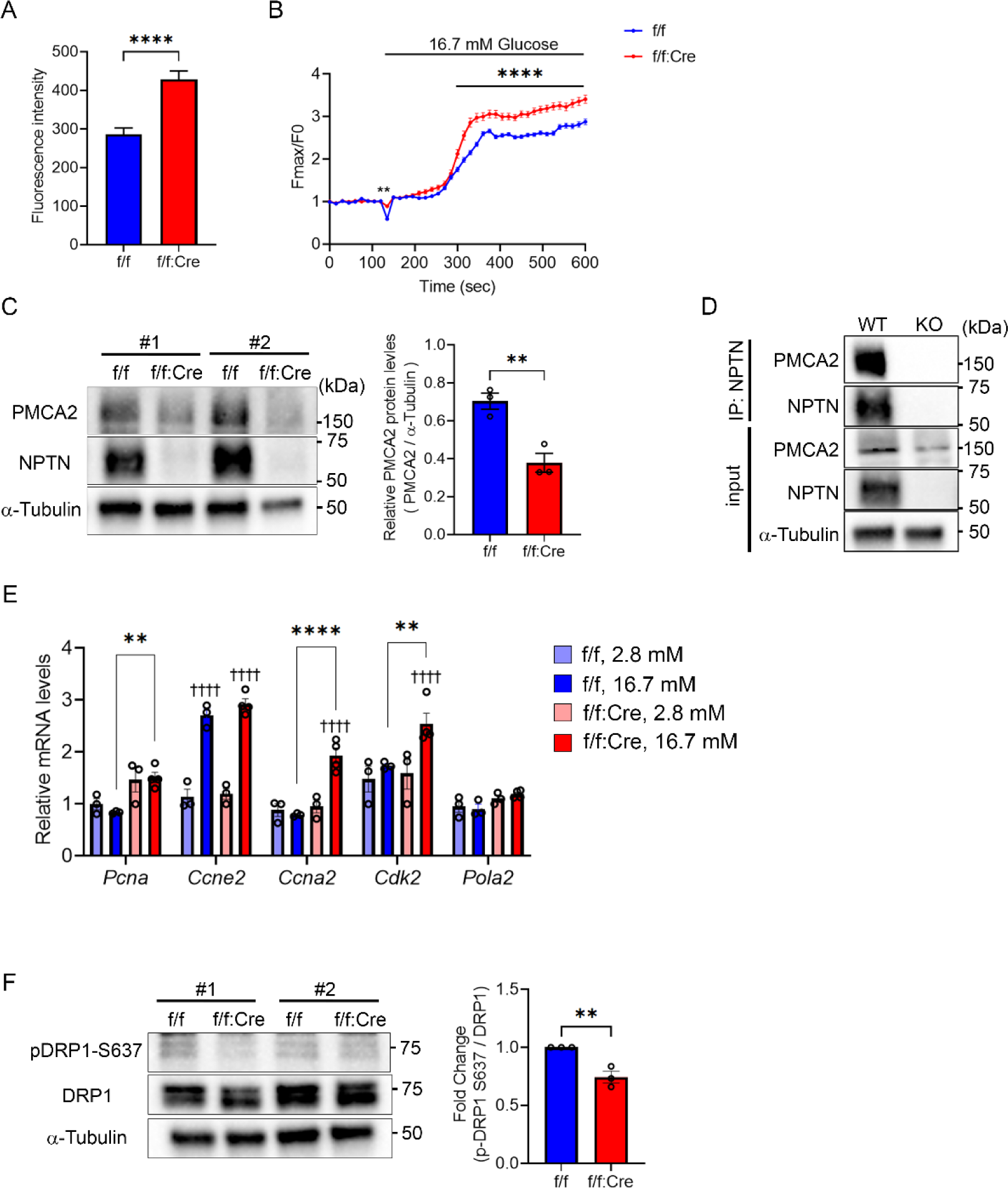
**NPTN stabilizes PMCA2 by the protein-protein interaction to regulate cytosolic Ca^2+^ homeostasis in βcells** (A) Basal cytosolic Ca^2+^ levels in islet cells from *Nptn*^f/f^ or *Nptn*^f/f^; *Ins1^Cre^* mice (*Nptn*^f/f^, n=73 cells, *Nptn*^f/f^; *Ins1^Cre^*, n=154 cells from 3 mice for each genotype. ****p<0.0001 by unpaired *t*-test). (B) Dynamic cytosolic Ca^2+^ levels in response to glucose stimulation in islet cells from *Nptn*^f/f^ or *Nptn*^f/f^; *Ins1^Cre^* mice (*Nptn*^f/f^, n=319 cells from 5 mice, *Nptn*^f/f^; *Ins1^Cre^*, n=136 cells from 4 mice. **p<0.01 and ****p<0.0001 by two-way ANOVA). (C) (Left) Representative blotting images of PMCA2, NPTN, and α-Tubulin in islets from *Nptn*^f/f^ or *Nptn*^f/f^; *Ins1^Cre^* mice. (Right) A quantification of PMCA2 protein levels normalized with α-Tubulin (*Nptn*^f/f^, n=3, *Nptn*^f/f^; *Ins1^Cre^*, n=3, **p<0.01 by unpaired *t*-test). (D) Representative blotting images of PMCA2, NPTN, and α-Tubulin in IP and input samples using *Nptn* WT or KO INS-1 832/13 cells. (E) qPCR analysis of NFATc2 target genes related to proliferation in *Nptn*^f/f^ or *Nptn*^f/f^; *Ins1^Cre^* islet cells cultured in the medium containing 2.8 mM or 16.7 mM glucose for 3 days. (*Nptn*^f/f^, 2.8 mM: n=3, *Nptn*^f/f^, 16.7 mM: n=3, *Nptn*^f/f^; *Ins1^Cre^*, 2.8 mM: n=3, *Nptn*^f/f^; *Ins1^Cre^*, 16.7 mM: n=4, †††† p<0.0001 by two-way ANOVA compared to 2.8mM for each genotype, **p<0.01 and ****p<0.0001 by two-way ANOVA). (F) (Left) Representative blotting images of pDRP1-S637, DRP1, and α-Tubulin in islets from *Nptn*^f/f^ or *Nptn*^f/f^; *Ins1^Cre^* mice. (Right) A quantification pDRP1-S637 protein levels normalized with DRP1 (*Nptn*^f/f^, n=3, *Nptn*^f/f^; *Ins1^Cre^*, n=3, **p<0.01 by unpaired *t*-test). f/f: *Nptn*^f/f^, f/f:Cre: *Nptn*^f/f^; *Ins1^Cre^*, ns: not statistically significant.

### NPTN transmits pro-inflammatory signal via TRAF6-NF-κB axis

We and the other research groups have previously reported that NPTN is involved in pro-inflammation by activating NF-κB signaling (20, 54). RNA-seq analysis with *Nptn*^f/f^; *Ins1^Cre^* islets showed lower expression of Reactome and KEGG pathways pertaining to several inflammatory signals (Figure 4A). Overexpression of *Nptn* in INS-1 832/13 cells significantly increased mRNA levels of pro-inflammatory cytokine and chemokine genes, which are NF-κB targets (55, 56) (Figure 4B). NF-κB signaling is activated by numerous discrete stimuli such as IL-1β and IFNγ that contribute to β cell failure in type 1 diabetes mellitus (T1D) (57, 58). Appropriately, NF-κB signaling was significantly activated after treating WT and *Nptn* KO INS-1 832/13 cells with cytokine mix (5 ng/ml IL-1β + 100 ng/ml IFNγ), as indicated by the rapid degradation of IκBα at 30 min after the treatment (59) (Figure 4C). However, the recovery of IκBα protein level was faster in *Nptn* KO INS-1 832/13 cells, suggesting that the NF-κB signaling is attenuated more quickly in *Nptn* KO cells. Similarly, the transcriptional induction of the NF-κB target genes by cytokine mix stimulation was inhibited in *Nptn* KO cells as compared to controls (Figure 4D). These results suggested that NPTN contributes to pro-inflammation via NF-κB signaling in pancreatic β cells. The crosstalk between β cells and immune cells in islets plays an important role in the progression of insulitis and β cell failure (60, 61). Given that NPTN is involved in the inflammation of β cells, NPTN could be a substantial contributor of the crosstalk. To verify this idea, we investigated if β cell-specific NPTN deficiency suppresses the development of diabetic phenotypes induced by multiple low dose-streptozotocin (MLD-STZ) injections, which is commonly used experimentally to produce a rodent model of T1D (62, 63). STZ injections increased non-fasting blood glucose levels over the weeks after the injections for the first 5 days in both *Nptn*^f/f^ and *Nptn*^f/f^; *Ins1^Cre^* mice (Figure 4E). However, the rate of increase was lower in *Nptn*^f/f^; *Ins1^Cre^* mice injected with STZ (Figure 4E). *Nptn*^f/f^; *Ins1^Cre^* mice also showed resistance to the development of glucose intolerance (Figure 4F). Moreover, β cell mass and serum insulin levels after glucose injection were higher in *Nptn*^f/f^; *Ins1^Cre^* mice injected with STZ as compared to controls (Figure 4G and H). Chemokines produced in β cells recruit circulating monocytes into islets, where they differentiate to macrophages (60, 61). Consistent with milder STZ-induced diabetic phenotypes of *Nptn*^f/f^; *Ins1^Cre^* mice, staining area of pan-macrophage marker, IBA1, was reduced in pancreatic sections of *Nptn*^f/f^; *Ins1^Cre^* mice injected with STZ (Figure 4I), indicating that NPTN contributes to islet inflammation and the development of insulitis and diabetes.

**Figure 4.**
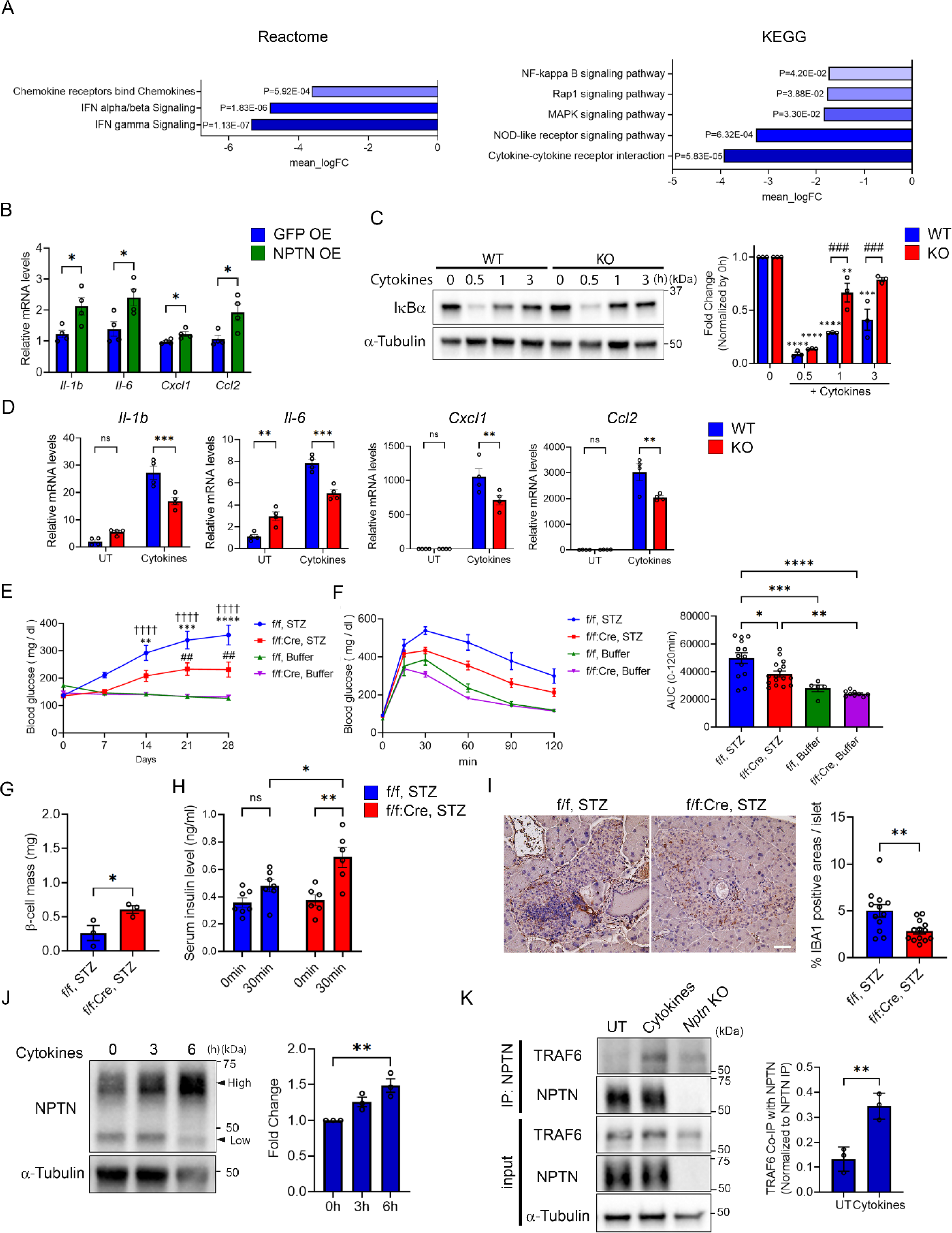
**NPTN deficiency inhibits islet inflammation induced by cytokines via TRAF6-NFκB axis** (A) Reactome and KEGG enrichment analyses for pathways downregulated in *Nptn*^f/f^; *Ins1^Cre^* islets (*Nptn*^f/f^: n=4, *Nptn*^f/f^; *Ins1^Cre^*: n=4). (B) qPCR analysis of NF-κB target genes related to pro-inflammatory cytokines in INS-1 832/13 cells with or without *Nptn* overexpression (n=4, *p<0.05 by unpaired *t-*test). (C) (Left) Representative blotting images of IκBα and α-Tubulin in *Nptn* WT or KO INS-1 832/13 cells treated with cytokine mix (5 ng/ml IL-1β + 100 ng/ml IFNγ) for indicated times. (Right) Fold changes of IkBα protein levels compared to 0h. The protein levels were normalized to α-Tubulin (n=3, **p<0.01, ***p<0.001, and ****p<0.0001 by two-way ANOVA compared to 0h for each genotype, ###p<0.001 by two-way ANOVA). (D) qPCR analysis of NF-κB target genes related to pro-inflammatory cytokines in *Nptn* WT or KO INS-1 832/13 treated with or without cytokine mix for 6 hours (n=4, **p<0.01 and ***p<0.001 by two-way ANOVA). (E) Non-fasting blood glucose levels in *Nptn*^f/f^ or *Nptn*^f/f^; *Ins1^Cre^* mice injected with STZ or buffer (*Nptn*^f/f^, STZ, n=16, *Nptn*^f/f^; *Ins1^Cre^*, STZ, n=16, *Nptn*^f/f^, Buffer, n=7, *Nptn*^f/f^; *Ins1^Cre^*, Buffer, n=10, **p<0.01, ***p<0.001 and ****p<0.0001 by two-way ANOVA compared to *Nptn*^f/f^; *Ins1^Cre^*, STZ, † † † † p<0.0001 by two-way ANOVA compared to *Nptn*^f/f^, Buffer, ##p<0.01 by two-way ANOVA compared to *Nptn*^f/f^; *Ins1^Cre^*, Buffer). (F) (Left) IP-GTT performed 25 days after the first injection with STZ or buffer. (Right) AUCs of the IP-GTT (*Nptn*^f/f^, STZ, n=13, *Nptn*^f/f^; *Ins1^Cre^*, STZ, n=16, *Nptn*^f/f^, Buffer, n=5, *Nptn*^f/f^; *Ins1^Cre^*, Buffer, n=9, *p<0.05, **p<0.01, ***p<0.001 and ****p<0.0001 by one-way ANOVA). (G) β cell mass of pancreas in *Nptn*^f/f^ or *Nptn*^f/f^; *Ins1^Cre^* mice injected with STZ 28 days after the first injection (*Nptn*^f/f^, STZ, n=3, *Nptn*^f/f^; *Ins1^Cre^*, STZ, n=3, *p<0.05 by unpaired *t*-test). (H) Serum insulin levels following glucose injection in *Nptn*^f/f^ or *Nptn*^f/f^; *Ins1^Cre^* mice injected with STZ 28 days after the first injection (*Nptn*^f/f^, STZ, n=7, *Nptn*^f/f^; *Ins1^Cre^*, STZ, n=6, *p<0.05 and *p<0.01 by two-way ANOVA). (I) (Left) Representative immunohistochemistry images of pancreas in *Nptn*^f/f^ or *Nptn*^f/f^; *Ins1^Cre^* mice injected with STZ 28 days after the first injection. Scale bar: 50 μm (Right) A quantification of IBA1-positive are per islet (*Nptn*^f/f^, n=12 islets from 3 mice *Nptn*^f/f^; *Ins1^Cre^*, n=13 islets from 3 mice, **p<0.01 by unpaired *t*-test). (J) (Left) Representative blotting images of NPTN and α-Tubulin in INS-1 832/13 cells treated with cytokine mix for indicated times. (Right) A quantification of NPTN protein levels at high molecular weight normalized to α-Tubulin (n=3, **p<0.01 by one-way ANOVA). (K) Representative blotting images of TRAF6, NPTN, and α-Tubulin in IP and input samples using Nptn WT or KO INS-1 832/13 cells treated with or without cytokine mix for 6 hours. (Right) A quantification of TRAF6 protein levels precipitated with NPTN. The protein levels were normalized to precipitated NPTP protein levels (n=3, **p<0.01 by impaired *t*-test). f/f: *Nptn*^f/f^, f/f:Cre: *Nptn*^f/f^; *Ins1^Cre^*, ns: not statistically significant.

Next, we investigated how NPTN mediates pro-inflammatory signaling via NF-κB pathway. Interestingly, western blot (WB) revealed an NPTN form of higher molecular weight (MW) which was elevated following cytokine mix stimulation, whereas the lower MW NPTN form was reduced (Figure 4J and Figure S6A). NPTN protein is highly glycosylated, which occurs within ER and Golgi apparatus. WB using purified proteins existing at plasma membrane showed only the high MW form, identifying as NPTN proteins at plasma membrane (Figure S6B). On the other hand, *Nptn* mRNA levels were not altered by the cytokine mix stimulation (Figure S6C), which suggested that NPTN proteins are transported from ER to plasma membrane in response to cytokine stimuli. In the NF-κB signal cascade, TNF receptor-associated factors (TRAFs), particularly TRAF2 and TRAF6, are pivotal mediators (64-66). Interestingly, a recent study has reported that the cytoplasmic region of NPTN contains TRAF6 binding motif, PxExxZ, and demonstrated an interaction between overexpressed these proteins (67). Similarly, we also confirmed the interaction between endogenous NPTN and TRAF6 in INS-1 832/13 (Figure 4K). This interaction was increased by cytokine mix treatment, indicating that TRAF6 transmits the pro-inflammatory signal from NPTN at the cell surface to the NF-κB signaling cascade. The previous study reported S100A8 and A9 as ligands of NPTN leading to the activation of NF-κB signaling (54). We then tested whether these ligands increase the expression of NF-κB target genes. Transcriptional inductions of *Il-1β* and *Il-6* were observed only in INS-1 832/13 cells treated with S100A9, but the magnitude was much smaller than that of cells treated with cytokine mix (Figure S6D and E). These inductions were negated in *Nptn* KO INS-1 832/13 cells (Figure S6D and E). Taken together, NPTN is involved in pro-inflammation in β cells via TRAF6-mediated NF-κB signal.

### MANF antagonizes inflammation and increases insulin secretion through NPTN

We previously demonstrated NPTN is a binding partner of MANF at the cell surface and is involved in the molecular mechanisms by which MANF inhibits inflammation (20). Therefore, we finally tested if MANF treatment results in similar phenotypes to that observed in *Nptn* KO cells. Consistent with our previous study, co-treatment with MANF peptides inhibited the transcriptional induction of NF-κB target genes induced by cytokine mix treatment in INS-1 832/13 cells (Figure 5A), whereas the anti-inflammatory effect of MANF was negated in *Nptn* KO INS-1 832/13 cells (Figure S7A). Expectedly, the interaction between NPTN and TRAF6, which was increased by cytokine stimuli, was also reduced by MANF co-treatment (Figure 5B). On the other hand, the NPTN protein levels in INS-1 832/13 cells treated with cytokine mix were similar regardless of MANF treatment (Figure 5B, input), suggesting that MANF treatment does not affect the increase of NPTN proteins at plasma membrane induced by cytokine mix stimulation. In addition to anti-inflammatory effect, we found NPTN deficiency increases GSIS *in vivo* and *in vitro*. Consecutive treatment with MANF proteins for 5 days significantly increased GSIS without affecting insulin content in the islets from *Nptn*^f/f^ mice (Figure 5C and D), whereas no increase was observed in *Nptn*^f/f^; *Ins1^Cre^* islets (Figure S7B and C). In the characterization of *Nptn*^f/f^; *Ins1^Cre^* mice or islets, we concluded that improved metabolic phenotypes caused by NPTN deficiency are due to reduced PMCA2 protein levels, which alter basal and dynamic Ca^2+^ levels in β cells. Interestingly, MANF treatment for 5 days significantly reduced PMCA2 protein levels along with NPTN in the islets (Figure 5E), suggesting that MANF treatment leads to a similar state to that of *Nptn* KO in islets. In summary, MANF reduces inflammation but also increases GSIS via NPTN in islets.

**Figure 5.**
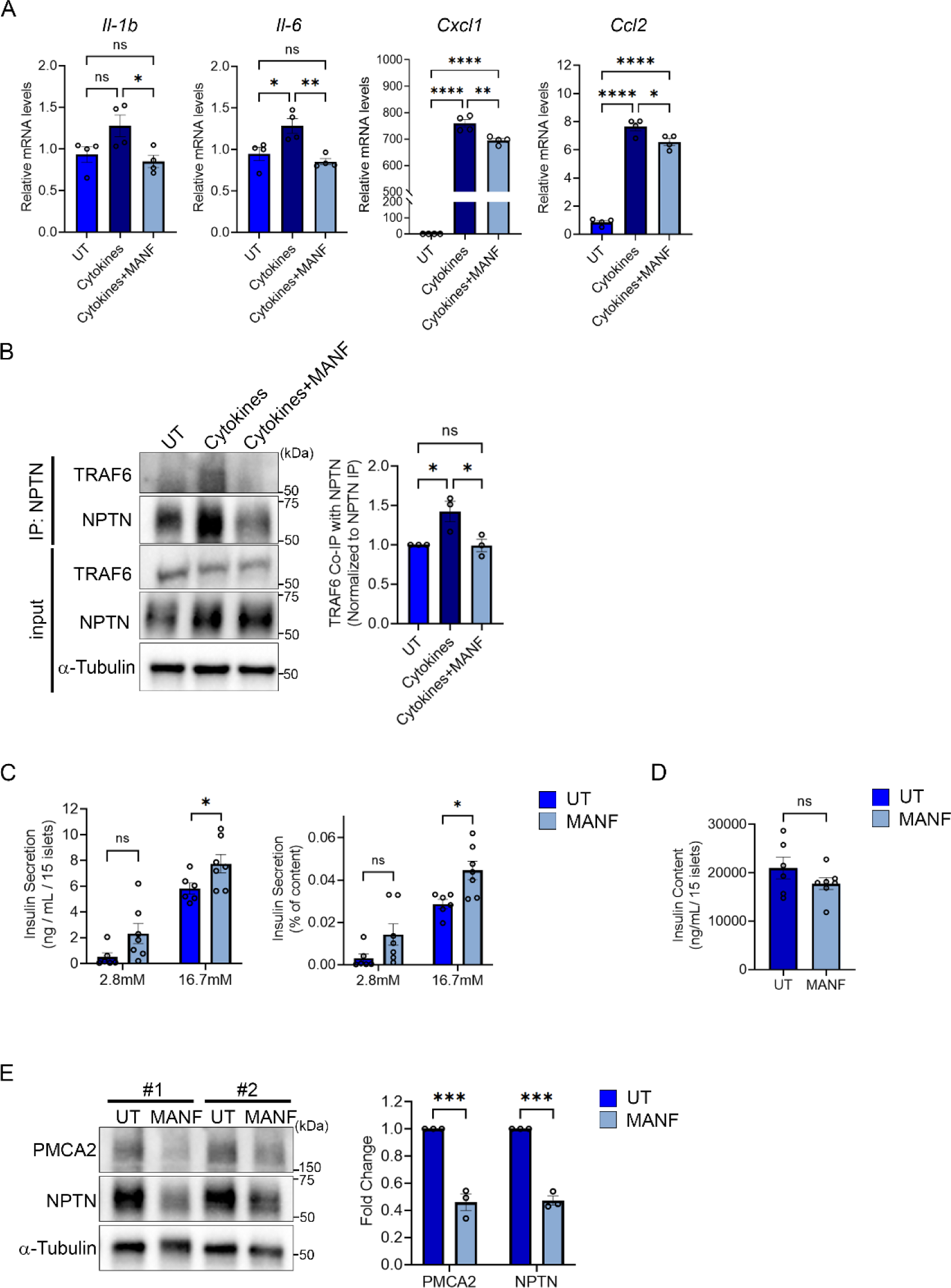
**MANF antagonizes NPTN roles in inflammation and insulin secretion** (A) qPCR analysis of NF-κB target genes related to pro-inflammatory cytokines in *Nptn* WT INS-1 832/13 treated with or without cytokine mix (5 ng/ml IL-1β + 100 ng/ml IFNγ) and MANF (5 μg/ml) for 6 hours (n=4, *p<0.05, **p<0.01 and ****p<0.0001 by one-way ANOVA). (B) Representative blotting images of TRAF6, NPTN, and α-Tubulin in IP and input samples using *Nptn* WT INS-1 832/13 cells treated with or without cytokine mix and MANF proteins for 6 hours. (Right) A quantification of TRAF6 protein levels precipitated with NPTN. The protein levels were normalized to precipitated NPTP protein levels (n=3, *p<0.05 by one-way ANOVA). (C) Static GSIS per islets (left) and normalized to insulin content (right) in primary islets form *Nptn*^f/f^ mice treated with or without MANF for 5 days (UT: n=6, *Nptn*^f/f^; MANF: n=7, *p<0.05 by unpaired *t*-test). (D) Insulin content in the islets used in (C). (E) (Left) Representative blotting images of PMCA2, NPTN, and α-Tubulin in islets from *Nptn*^f/f^ or *Nptn*^f/f^; *Ins1^Cre^* mice treated with or without MANF for 5 days (Right) A quantification of PMCA2 and NPTN protein levels normalized with α-Tubulin (*Nptn*^f/f^, n=3, *Nptn*^f/f^; *Ins1^Cre^*, n=3, ***p<0.001 by unpaired *t*-test). UT: untreated, f/f: *Nptn*^f/f^, f/f:Cre: *Nptn*^f/f^; *Ins1^Cre^*, ns: not statistically significant.

## Discussion

To understand the role of NPTN in pancreatic β cells, we conducted metabolic characterization of β cell-specific *Nptn* KO mice and observed improved glucose tolerance as well as increased glucose-stimulated insulin secretion, proliferation, and mitochondrial number in β cells. NPTN stabilizes PMCA2 protein through protein-protein interaction, which is crucial for regulating cytosolic Ca^2+^ homeostasis. The improved metabolic phenotypes in β cell-specific *Nptn* KO mice are due to increased basal and dynamic Ca^2+^ levels caused by altered PMCA2 protein levels in β cells. In addition to β cell functions, NPTN is involved in pro-inflammation by activating the TRAF6-NF-κB axis. The NPTN-mediated inflammation in β cells contributes to the development of STZ-induced diabetic phenotypes. We also demonstrated that the secretory protein MANF antagonizes these NPTN-mediated physiologies in β cells, and long-term MANF treatment affected both NPTN and PMCA2 protein levels.

NPTN is originally identified as a synaptic glycoprotein (23). Accordingly, most of earlier studies mainly focused on its physiological or disease-relevant roles in nervous systems. To date, it has been reported that NPTN has various roles in neurons such as synaptogenesis, neurite outgrowth, proper long-term synaptic plasticity, and transportation of lactate (22, 25, 45, 67-71). These roles are determined by the binding partner of NPTN and biochemical analyses suggested that NPTN proteins are interacted with themselves in the form of cis-or trans-homophilic binding and a variety of other proteins including PMCAs, fibroblast growth factor receptors (FGFRs), GABA_A_ receptors, monocarboxylate transporter 2 (MCT2) and some cellular signal mediators such as TRAF2/6 and GRB2 (22, 25, 44-46, 54, 67-71). Along with NPTN, some of these binding partners are expressed not only in nervous systems but also in peripheral tissues, implying that NPTN may have important roles regulating physiological and pathophysiological functions in non-neuronal cells by the similar molecular mechanisms. Indeed, we demonstrated that NPTN is involved in regulating proper insulin secretion, proliferation, and mitochondria number in β cells through the interaction with PMCA2 protein. Also, NPTN transmits inflammatory signals by the binding of TRAF6, which contributes to the development of diabetic phenotypes. Furthermore, other studies suggested that NPTN regulates T cell activation and proliferation of keratinocytes by interacting with PMCAs and TRAF2, respectively (44, 54).

We demonstrated that NPTN is interacted with PMCA2 and this interaction is important for the stabilization of PMCA2 protein and the functional capacities of β cells. Interestingly, heterozygous inactivation of PMCA2 in mice increases GSIS, proliferation, and mass in β cells (72). These phenotypes are also observed in β cell-specific *Nptn* KO mice, supporting our conclusion that the interaction between NPTN and PMCA2 proteins plays a pivotal role in regulating β cell physiology. It is well known that cytosolic Ca^2+^ is an important second messenger responsible for the transmission of physiological and pathophysiological signals modulating various cell processes. To more define NPTN roles, further investigations in the other Ca^2+^ sensitive tissues such as muscle and kidney could be intriguing.

It was shown that NPTN recruits TRAF6 to provoke NF-kB signaling in response to inflammatory stimuli. However, a ligand specific to NPTN has not been identified. According to genome-wide sequence searches, NTPN is most closely related to the other Ig superfamily member, CD147 (22). Interestingly, CD147 was identified as another component of a protein complex with NPTN and PMCAs, which is required for stabilization of PMCA2 protein (45). The role of CD147 in the inflammation was well-studied and A100A8/9 and CycrophilinA are shown as ligands activating NF-kB signaling via CD147 (73, 74). Sakaguchi et al. has demonstrated that S100A8 and S100A9 are bound with Np65 and Np55, respectively, to induce the expression of NF-κB target genes (54). We also confirmed an increase of cytokine gene expression in INS-1 832/13 cells treated with S100A9, whereas those expression was similar regardless of the treatment in *Nptn* KO cells. However, the magnitude of transcriptional induction by S100A9 was much smaller than that induced by cytokine mix stimulation, indicating it is uncertain that S100A9 is a specific ligand for NPTN in β cells. As an alternative molecular mechanism by which NPTN elicits inflammatory signal, NPTN may form hetero dimer/oligomeric receptor complex with cytokine receptors in the similar way to the case of PMCAs, which could modulate the inflammatory signals derived from these receptors. Meanwhile, we found an increase of NPTN proteins at plasma membrane in response to cytokine mix stimulation. NPTN is also known as an adhesion molecule and is important for synaptogenesis (21, 22, 67). For decades, adhesion molecules have been suggested to be involved in inflammation (75, 76). Of note, recently it was reported that numbers of cell adhesion molecule 1 (CADM1)-positive islet endocrine and myeloid cells adjacent to CD8^+^ T cells, which are well known to destroy insulin-producing β cells (77), are increased in individuals with T1D (78). CADM-1 is an Ig domain-containing plasma membrane protein that mediates homotypic and heterotypic cell-to-cell contact (79, 80). Remarkably, the suppression of *Cadm1* promotes insulin secretion and β cell mass (81-83) similar to the phenotypes we observed in β cell-specific *Nptn* KO mice. β cell-specific KO mice are significantly resistant to development of STZ-induced diabetic phenotypes, therefore, NPTN could mediate cell-cell contact between islet cells and immune cells in addition to recruitment of immune cells into islets by increasing chemokine expression.

Several studies revealed effects of MANF on glucose homeostasis (4). In this study, we observed an increase of GSIS in primary islets treated with MANF proteins for 5 days, which was negated in *Nptn* KO islets. However, our previous examination did not admit a significant effect of a short-term MANF treatment (24 hours) on insulin secretion (5). MANF is known to promote β cell proliferation (13, 14) and it has been demonstrated that a consecutive treatment of whole islets with MANF proteins for 5 days enhances the proliferation rate of human and murine β cells (5, 14). Hence, it is suggested that our observation of increased GSIS is due to an increased β cell mass and MANF treatment for 24 hours could be too short to increase enough number of β cells for improving GSIS. On the other hand, we confirmed anti-inflammatory effect in INS-1 832/13 cells treated with MANF proteins for 8 hours. Given that MANF effects on GSIS and anti-inflammation were negated in *Nptn* KO islets/cells, both of the MANF effects are mediated via NPTN. However, the mechanisms how MANF suppresses NPTN roles in GSIS and inflammation could be different. Of note, islets treated with MANF proteins for 5 days exhibited decreased NPTN protein levels, whereas no change was observed in INS-1 832/13 cells treated for 8 hours. In addition to the signal transduction of MANF mediated by NPTN at plasma membrane, it is suggested that MANF is internalized through the N-terminal saposin-like domain and transferred into ER (19). Also, it is shown that MANF existing in ER stabilizes certain BiP-client complex, which may enhance the efficiency of client transfer to downstream ER quality control effectors including ER-associated degradation (ERAD) (18). Thus, as a putative mechanism, MANF might be involved in facilitating the handover of NTPN in ER to ERAD leading to the decrease in the protein levels of NPTN.

This study unveiled NPTN roles in β cells, which are important for mitigating insulin secretion and promoting islet inflammation. These roles are inhibited by MANF treatment, suggesting that MANF exerts insulin secretory and anti-inflammatory effects through binding to NPTN. Thus, inhibiting NPTN function or expression could be a good strategy to prevent or reverse β cell failure or loss in diabetes.

## Methods

### Animals

All animal experiments were performed according to procedures approved by the Institutional Animal Care and Use Committee at the Washington University School of Medicine (20-0334). *Ins1^Cre^* mice are obtained from Jackson Laboratory (RRID:IMSR_JAX:026801). *Ntpn*^flox/flox^ mice are generated by the Genome Engineering and iPSC Center (GEiC) at Washington University in St. Louis.

### *in vivo* physiology and pancreatic insulin content

Intraperitoneal glucose tolerance test (IP-GTT), intraperitoneal insulin tolerance test (IP-ITT) and *in vivo* glucose-stimulated insulin secretion test were performed according to standard procedures of the NIH-sponsored National Mouse Metabolic Phenotyping Centers (http://www.mmpc.org). To measure insulin content in whole pancreas, excised pancreata were weighed and homogenized in ice-cold acid ethanol (1.5% 12 N HCl in 70% ethanol). Homogenized pancreata were incubated at −20°C for 48 hours. Pancreatic and serum insulin content were measured by Ultra Sensitive Mouse Insulin ELISA Kit (Crystal Chem; 90080). 7 weeks old mice were used in the analyses except for that with STZ-injected mice.

### Islet isolation and dispersion

Murine islets were isolated from the mice at 7-12 weeks old (7 weeks old: RNA-seq and 8-12 weeks old: the other analyses) as described previously (84). After isolation, islets were incubated overnight in RPMI 1640 medium (Gibco, 11875085) (11.1mM glucose, 10% PBS, and Penicillin-Streptomycin (Gibco, 15140122) to allow them to recover from the digestion damages. For MANF treatment experiments, the islets were cultured for 5 days in RPMI 1640 medium (11.1 mM glucose, 10% FBS and Penicillin-Streptomycin) with or without 5 μg/ml recombinant human MANF protein (R&D systems, 3748-MN). For EdU cell proliferation assay and Ca^2+^ measurement, 50-70 islets were collected in a 1.5 ml tube for each well of 8-well chamber. The islets were incubated in 200 μl Accutase solution (SIGMA, A6964) at 37 ℃ for 6 min after washing them with HBSS (Gibco, 14175145). The islets were dissociated by pipetting gently followed by adding 400 μl RPMI 1640 medium. After centrifuge at 300 x g, for 4min, the pelleted islet cells were resuspended with RPMI 1640 medium and plated on a laminin-coated well of 8-well chamber.

### Static glucose-stimulated insulin secretion (GSIS) assay

15 islets were equilibrated in Krebs-Ringer bicarbonate Hepes (KRBH) buffer (128.8 mM NaCl, 4.8 mM KCl, 1.2 mM KH_2_PO_4_, 1.2 mM MgSO_4_, 2.5 mM CaCl_2_, 20 mM HEPES, 5 mM NaHCO3, 0.1% BSA, pH7.4) at 2.8 mM glucose for 1 hour at 37 °C. Then, the islets were incubated in new KRBH buffer at 2.8 mM glucose for 30 min followed by 11 mM glucose for 30 min. Supernatant was collected after the incubation in each solution. At the end of assay, the islets were lysed overnight in acid-ethanol. Secreted insulin and insulin content were measured by Ultra Sensitive Mouse Insulin ELISA Kit.

### Immunofluorescence and measurement of β cell mass

For β cell mass study, pancreata were weighed and fixed in 4% PFA overnight at 4℃ and praffin-embedded for sectioning. 4 groups of sections with 5 μm thickness were made 75 μm apart between groups. After rehydration and antigen retrieval in 10 mM sodium citrate buffer (pH 6.0), sections were permeabilized in 0.3% Triton-X prior to blocking in 2% BSA. Primary antibodies were treated overnight at 4℃. Incubation with secondary antibodies and DAPI was for 1 hour at RT. Slides were mounted with ProLong Diamond Antifade Mountant (Invitrogen, P36970). For the other saining, pancreata were perfused with 4% PFA and fixed overnight at 4℃. After replacement with 30% sucrose gradually, pancreata were frozen in Tissue-Tek® O.C.T. Compound (Sakura FineTek, 4583). Frozen pancreata were sectioned with 10 μm thickness. The procedure after permeabilization was performed in the same way as insulin staining. β cell mass (mg per pancreas) was calculated by multiplying relative insulin-positive area (the percentage of insulin positive area over total pancreas area) by pancreas weight. 7 weeks old mice were used in the analyses. All Images analyses were performed with image J. Antibody details can be found in Table S1.

### EdU cell proliferation assay

Dispersed islet cells were cultured in RPMI 1640 medium containing either 2.8 mM or 16.7 mM glucose suppled with 10% FBS and Penicillin-Streptomycin for 3 days. The half of medium was changed every 24 hours. 10 μM EdU was added in the culture medium for the last 24 hours of stimulation. Incorporated EdU was detected by Click-iT EdU Alexa Fluor 488 Imaging Kit (Invitrogen, C10337). To determine β cells, the cells were stained with insulin antibody and DAPI following EdU detection by a general immunofluorescence protocol. The areas for imaging were picked up randomly and the images were analyzed with Image J. Antibody details can be found in Table S1.

### Ca^2+^ measurement

Dispersed islet cells were laded with 4 μm Calbryte 520 AM (AAT Bioquest 20650) in loading solution (2 mM Glucose in KRBH buffer) at 37℃ for 1 hour. After washing the cells with loading solution, 8-well chamber was inserted on the stage of microscope with incubation system (37℃, 5% CO_2_). First, the basal cytosolic Ca^2+^ levels were measured every 15 sec for 2 min. After the solution was changed to KRBH buffer containing 16.7 mM glucose, the dynamic changes of cytosolic Ca^2+^ levels were measured every 15 sec up to 10 min in total. The fluorescent emission signals were collected using Zeiss LSM 880 Airyscan two-photon microscope. Images were analyzed with image J.

### Cells and cytokine treatment

INS-1 832/13 cells were cultured in RPMI 1640 medium (11.1 mM glucose, 10% FBS, 1 mM sodium pyruvate (Gibco, 11360070), 2-mercaptoethanol (SIGMA, M6250), Penicillin-Streptomycin). *Nptn* WT ant KO INS-1 832/13 cells were generated by GEiC at Washington University in St. Louis. For cytokine treatments, the medium was changed to serum-free RPMI 1640 medium 1 hour prior to the treatment, cytokine treatment was performed with or without MANF by medium change. All recombinant proteins were obtained from R&D systems: IL-1β (R&D systems, 501-RL), IFNγ (R&D systems, 585-IF), S100A8 (R&D systems, 9877-S8) and S100A9 (R&D systems, 2065-S9). Concentrations for each cytokine are specified in the figure legends.

### Statistical analysis

Statistical analysis was performed by unpaired and paired t tests and one- and two-way ANOVA with Tukey’s or Dunnett’s tests. Statistical tests are specified in figure legends. P < 0.05 was considered statistically significant. Data are shown as means ± SEM unless otherwise noted.

Further methods details are available in the Supplemental Appendix.

## Author contributions

RAK and FU conceived the experimental design. RAK performed all experiments. FU supervised the data analyses. RAK, DH, and FU wrote the manuscript. All authors edited and reviewed the manuscript.

## Acknowledgements

This work was partly supported by the grants from the National Institutes of Health (NIH)/NIDDK (DK132090, DK020579).

