## Supporting Information for "Dual Role of Neuroplastin in Pancreatic β Cells: Regulating Insulin Secretion and Promoting Islet Inflammation"

Supplemental Appendix

Extended Methods

Extended References 85-91

Figure S1. Pancreatic β cell-specific *Nptn* KO mice.

Figure S2. Metabolic characterization of β cell-specific *Nptn* KO female mice.

Figure S3. Additional gene set enrichment analysis on the islets.

Figure S4. The expression of β cell key genes in β cell-specific *Nptn* islets.

Figure S5. Additional analyses of the PMCA2 protein stability in NPTN-deficient cells.

Figure S6. Additional analyses of the NPTN roles in inflammation.

Figure S7. Additional analyses of MANF effects on the NPTN-mediated inflammation and insulin secretion.

Table S1. Antibody list used in this study.

Table S2. Primer list used in this study.

### Extended Methods

### RNA Sequencing and Analysis

Samples were prepared according to library kit manufacturer's protocol, indexed, pooled, and sequenced on an Illumina NovaSeq 6000. Basecalls and demultiplexing were performed with Illumina's bcl2fastq2 software. RNA-seq reads were then aligned and quantitated to the Ensembl release 101 primary assembly with an Illumina DRAGEN Bio-IT on-premise server running version 3.9.3-8 software.

All gene counts were then imported into the R/Bioconductor package EdgeR (85) and TMM normalization size factors were calculated to adjust for samples for differences in library size. Ribosomal genes and genes not expressed in the smallest group size minus one samples greater than one count-per-million were excluded from further analysis. The TMM size factors and the matrix of counts were then imported into the R/Bioconductor package Limma (86). Weighted likelihoods based on the observed mean-variance relationship of every gene and sample were then calculated for all samples and the count matrix was transformed to moderated log 2 counts-per-million with Limma's voomWithQualityWeights (87). The performance of all genes was assessed with plots of the residual standard deviation of every gene to their average log-count with a robustly fitted trend line of the residuals. Differential expression analysis was then performed to analyze for differences between conditions and the results were filtered for only those genes with Benjamini-Hochberg false-discovery rate adjusted p-values less than or equal to 0.05.

For each contrast extracted with Limma, global perturbations in known Gene Ontology (GO) terms, MSigDb, and KEGG pathways were detected using the R/Bioconductor package GAGE (88) to test for changes in expression of the reported log 2 fold-changes reported by Limma in each term versus the background log 2 fold-changes of all genes found outside the respective term. The R/Bioconductor package heatmap3 (89) was used to display heatmaps across groups of samples for each GO or MSigDb term with a Benjamini-Hochberg false-discovery rate adjusted p-value less than or equal to 0.05. Perturbed KEGG pathways where the observed log 2 fold-changes of genes within the term were significantly perturbed in a single-direction versus background or in any direction compared to other genes within a given term with p-values less than or equal to 0.05 were rendered as annotated KEGG graphs with the R/Bioconductor package Pathview (90).

To find the most critical genes, the Limma voomWithQualityWeights transformed log 2 counts-per-million expression data was then analyzed via weighted gene correlation network analysis with the R/Bioconductor package WGCNA (27). Briefly, all genes were correlated across each other by Pearson correlations and clustered by expression similarity into unsigned modules using a power threshold empirically determined from the data. An eigengene was then created for each de novo cluster and its expression profile was then correlated across all coefficients of the model matrix. Because these clusters of genes were created by expression profile rather than known functional similarity, the clustered modules were given the names of random colors where grey is the only module that has any pre-existing definition of containing genes that do not cluster well with others. These de-novo clustered genes were then tested for functional enrichment of known GO terms with hypergeometric tests available in the R/Bioconductor package clusterProfiler (91). Significant terms with Benjamini-Hochberg adjusted p-values less than 0.05 were then collapsed by similarity into clusterProfiler category network plots to display the most significant terms for each module of hub genes in order to interpolate the function of each significant module. The information for all clustered genes for each module were then combined with their respective statistical significance results from Limma to determine whether or not those features were also found to be significantly differentially expressed.

**Electron microscopy**

Islets were fixed in a modified Karnovsky's fixative of 3% glutaraldehyde and 1% paraformaldehyde in 0.1M sodium cacodylate buffer and centrifuged at 1000 rpm to a pellet. Then, the islets were post-fixed in 2% osmium tetroxide in 0.1M sodium cacodylate buffer for 1 hour, en bloc stained with 3% aqueous uranyl acetate for 30 min, dehydrated in graded ethanols and embeded in PolyBed 812 (Polysciences, # 08792-1). Embedded Islet pellets were sectioned at 90 nm thick, post stained with Venable's lead citrate and viewed with a JEOL model 1400EX electron microscope (JEOL). Digital images were acquired using the AMT NanoSprint 12A-B (Advanced Microscopy Technology) CMOS, 12 megapixel TEM camera. Images were analyzed with Image J.

**Quantification of mitochondrial DNA contents**

Genome was extracted from islets using NucleoSpin Tissue (TaKaRa; 740952.5) and 10 ng genomic DNA was used for qPCR. Mitochondrial DNA copy numbers were normalized to nuclear DNA (34). Primers used for qPCR are listed in Supplemental Table S2.

**Nucleofection**

$4x10^6$ INS-1 832/13 cells were introduced with 2 μg pmax-GFP or Np55-GFP plasmids using P3 Primary Cell 4D-Nucleofector X kit (Lonza, V4XP-3024) according to the manufacturer's protocol. pmax-GFP was provided along with the kit. The details of Np55-GFP were described previously (20). The cells were used 48 hours after the nucleofection.

**Real-time qPCR**

Total RNA was isolated using RNeasy Kits (Qiagen, 74106) and cDNA libraries were generated using high-capacity cDNA reverse transcription kits (Applied Biosystems, 4368814). Relative amounts of each transcript were calculated by the ΔΔCt method and normalized to human 18S rRNA. Quantitative PCR was performed with the Applied Biosystems ViiA7 using PowerUp™ SYBR™ Green Master Mix (Applied Biosystems, A25741). Primers used for qPCR are listed in Supplemental Table S2.

**Western blotting and Purification of cell surface proteins**

For detecting NPTN and PMCA2, protein was extracted using 1% Digitonin lysis buffer (20 mM Tris, 50 mM NaCl, 2 mM $MgCl_2$). For pDrp1-S634, Drp1 and IκBα, protein was extracted using M-PER™ Mammalian Protein Extraction Reagent (Thermo; 78501). Both lysis buffers were supplied with protease inhibitor cocktail (Roche, 11697498001) and phosphatase inhibitor cocktail (Roche, 04906845001). Cell surface proteins were biotinylated with 0.25 mg/ml EZ-Link Sulfo-NHS-SS-Biotin in PBS and purified according to the manufacturer's protocol of Pierce™ Cell Surface Protein Isolation Kit (Thermo, 89881). An equivalent amount of total protein was loaded onto the SDS-polyacrylamide gel. Proteins were probed with primary and corresponding secondary antibodies. Antibody details can be found in Supplemental Table S1.

**Multiple low dose streptozotocin (MLD-STZ) injections**

Male mice at 8 weeks old received intraperitoneally 40 mg/kg STZ (SIGMA, S0130) dissolved in citrate buffer on pH 4.5 for 5 consecutive days. The daily dose of STZ was determined by the daily weight measurement of each mouse. The injection of the substance was performed within 5 min after STZ solution was prepared.

**Immunohistochemistry**

Paraffin sections with 5 μm thickness were subject to antigen retrieval in 10 mM sodium citrate buffer (pH 6.0) and permeabilized in 0.1% Triton-X after rehydration. Endogenous peroxidase was quenched with BLOXALL® Endogenous Blocking Solution (Vector Laboratories, SP-6000). Sections were blocked in 2% BSA followed by treated with primary IBA1 antibodies overnight at 4°C. IBA1 signals were detected by ImmPRESS® HRP Horse Anti-Goat IgG Polymer Detection Kit (Vector Laboratories, MP-7405) and DAB Substrate Kit (Vector Laboratories, SK-4100). For the counterstain of nuclei, hematoxylin staining was performed using Hematoxylin QS Counterstain (Vector Laboratories, H-3404). Slides were mounted with VectaMount® Permanent Mounting Medium (Vector Laboratories, H-5000-60). Antibody details can be found in Supplemental Table S1.

### Immunoprecipitation

For precipitating NPTN and PMCA2, cells were lysed in 1% Digitonin buffer (20 mM Tris, 50 mM NaCl, 2 mM $MgCl_2$). For NPTN and TRAF6, cells were lysed in RIPA buffer (20 mM Tris, 100 mM NaCl, 1 mM EDTA, 10% Glycerol, 0.1% SDS, 1% Triton-X, 1 mM N-Ethylmaleimide (NEM)). Both lysis buffers were supplied with protease inhibitor cocktail and phosphatase inhibitor cocktail. Equivalent proteins were incubated with 1 μg NPTN antibodies overnight at 4°C after pre-cleaning. Protein G Sepharose 4B (Invitrogen, 101241) were added for 3 h at 4 °C. Beads were washed three times with washing buffer (20 mM Tris, 150 mM NaCl, 0.5% Digitonin) followed by a rinse with washing buffer without Digitonin (20 mM Tris, 150 mM NaCl). Samples were subject to western blot. Antibody details can be found in Supplemental Table S1.

**Extended References**

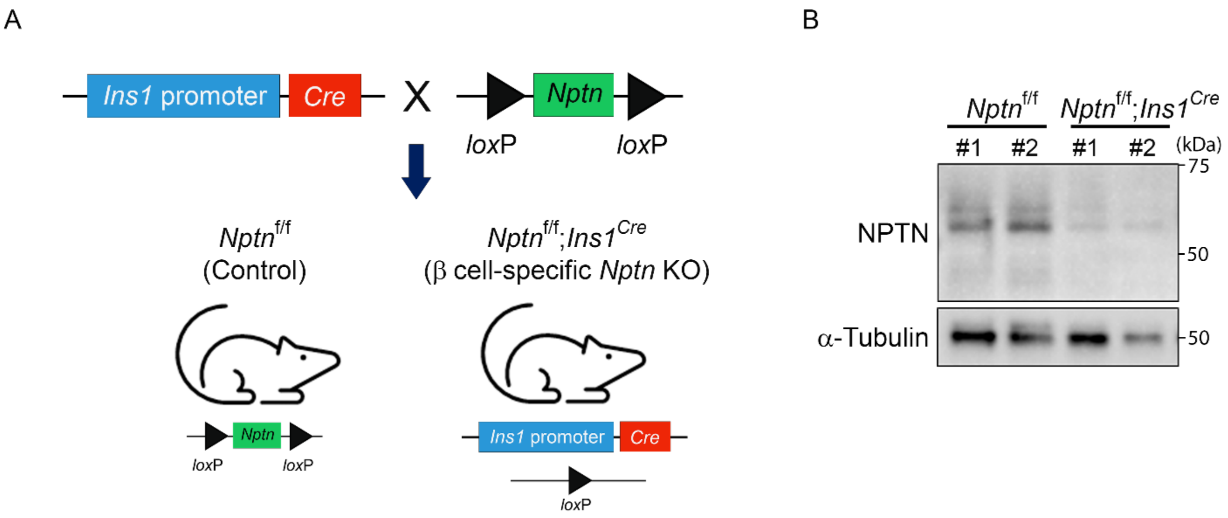


**Figure S1. Pancreatic β cell-specific *Nptn* KO mice.**

(A) A schematic of generation of β cell-specific *Nptn* KO mice. (B) Representative blotting images of NPTN and α-Tubulin in islets from *Nptn*$^{f/f}$ or *Nptn*$^{f/f}$; *Ins1*$^{Cre}$.

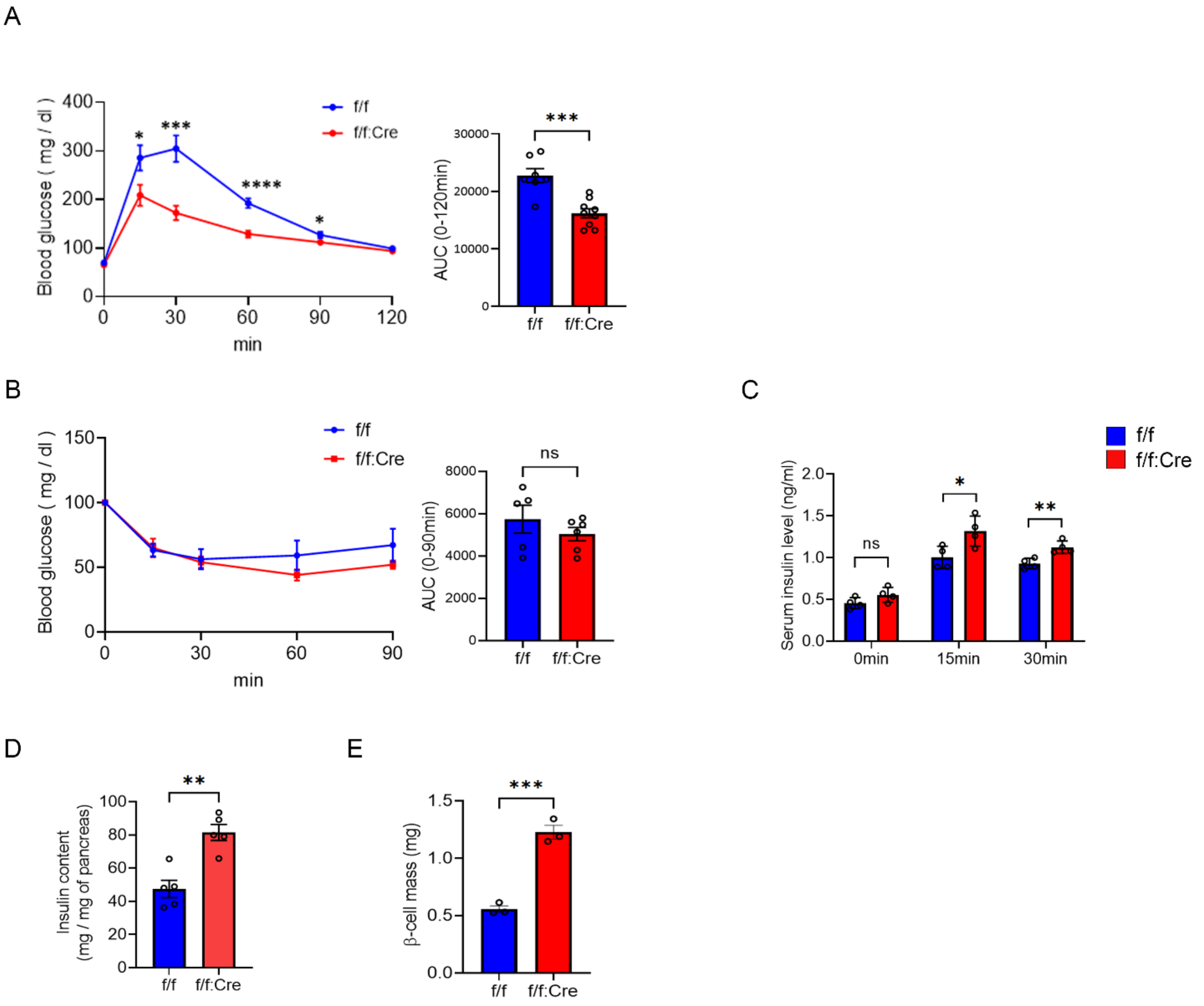


**Figure S2. Metabolic characterization of β cell-specific *Nptn* KO female mice.**

(A) (Left) IP-GTT with *Nptn*$^{f/f}$ or *Nptn*$^{f/f}$; *Ins1*$^{Cre}$ female mice. (Right) AUCs of the IP-GTT (*Nptn*$^{f/f}$: n=7, *Nptn*$^{f/f}$; *Ins1*$^{Cre}$: n=9, *p<0.05, ***p<0.001, and ****p<0.0001 by unpaired *t*-test). (B) (Left) IP-ITT with *Nptn*$^{f/f}$ or *Nptn*$^{f/f}$; *Ins1*$^{Cre}$ female mice. (Right) AUCs of the IP-ITT (*Nptn*$^{f/f}$: n=5, *Nptn*$^{f/f}$; *Ins1*$^{Cre}$: n=6). (C) Serum insulin levels following glucose injection in *Nptn*$^{f/f}$ or *Nptn*$^{f/f}$; *Ins1*$^{Cre}$ female mice (*Nptn*$^{f/f}$: n=4, *Nptn*$^{f/f}$; *Ins1*$^{Cre}$: n=4, *p<0.05 and **p<0.01 by unpaired *t*-test). (D) Insulin content of whole pancreas from *Nptn*$^{f/f}$ or *Nptn*$^{f/f}$; *Ins1*$^{Cre}$ female mice (*Nptn*$^{f/f}$: n=5, *Nptn*$^{f/f}$; *Ins1*$^{Cre}$: n=5, **p<0.01 by unpaired *t*-test). (E) β cell mass of pancreas in *Nptn*$^{f/f}$ or *Nptn*$^{f/f}$; *Ins1*$^{Cre}$ female mice (*Nptn*$^{f/f}$: n=3, *Nptn*$^{f/f}$; *Ins1*$^{Cre}$: n=3, ***p<0.001 by unpaired *t*-test). f/f: *Nptn*$^{f/f}$, f/f:Cre: *Nptn*$^{f/f}$; *Ins1*$^{Cre}$, ns: not statistically significant.

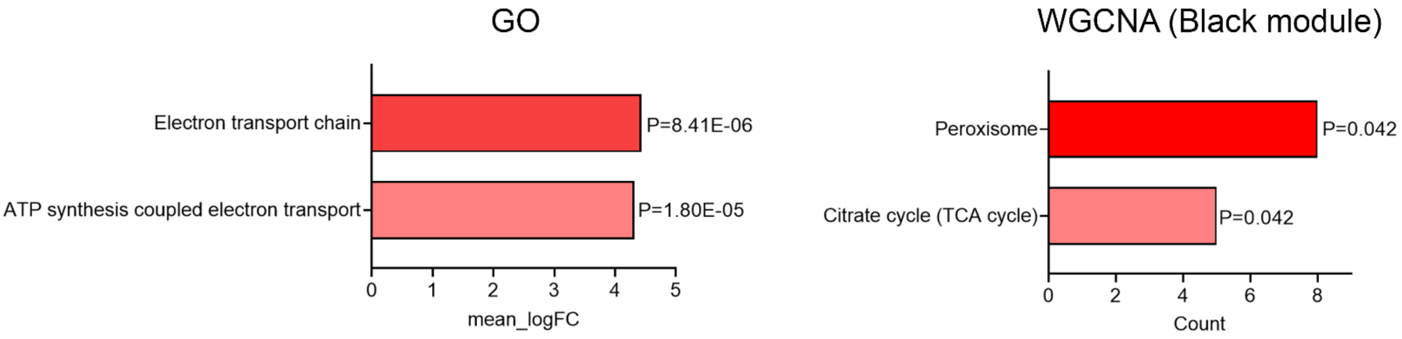


**Figure S3. Additional gene set enrichment analysis on the islets.**

(Left) GO enrichment analysis and for gene sets upregulated in *Nptn*$^{f/f}$; *Ins1*$^{Cre}$ islets and (Right) KEGG pathway of WGCNA black module (*Nptn*$^{f/f}$: n=4, *Nptn*$^{f/f}$; *Ins1*$^{Cre}$: n=4).

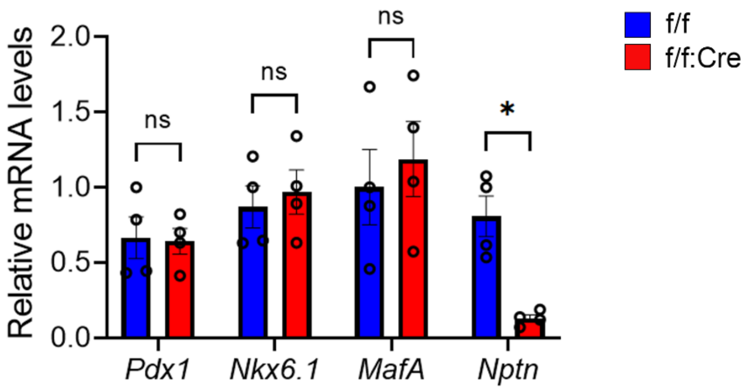


**Figure S4. The expression of β cell key genes in β cell-specific *Nptn* islets.**

The mRNA levels of β cell key transcription factors were similar in islets regardless of genotypes. (*Nptn*$^{f/f}$: n=4, *Nptn*$^{f/f}$; *Ins1*$^{Cre}$: n=4, *p<0.05 by unpaired *t*-test). f/f: *Nptn*$^{f/f}$, f/f:Cre: *Nptn*$^{f/f}$; *Ins1*$^{Cre}$, ns: not statistically significant.

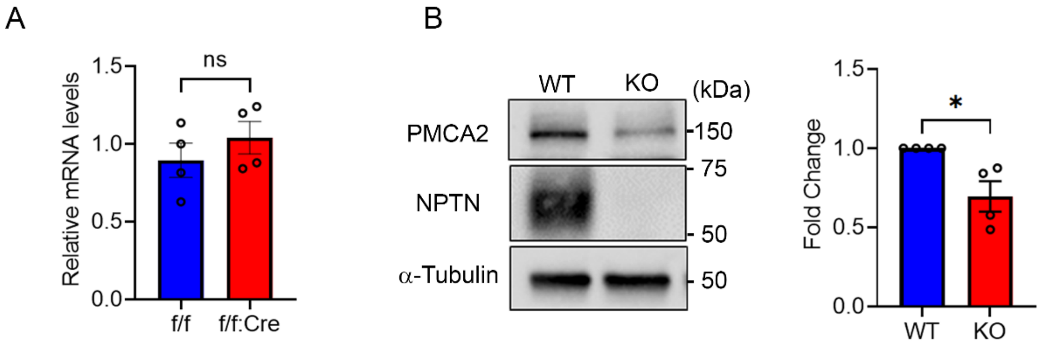


**Figure S5. Additional analyses of the PMCA2 protein stability in NPTN-deficient cells.**

(A) *Pmca2* mRNA levels were similar in islets regardless of genotypes (*Nptn*$^{f/f}$: n=4, *Nptn*$^{f/f}$; *Ins1*$^{Cre}$: n=4,). (B) (Left) Representative blotting images of PMCA2, NPTN, and α-Tubulin in *Nptn* WT or KO INS-1 832/13 cells. (Right) A quantification of PMCA2 protein levels normalized to α-Tubulin (WT: n=4, KO: n=4, *p<0.05 by unpaired *t*-test). f/f: *Nptn*$^{f/f}$, f/f:Cre: *Nptn*$^{f/f}$; *Ins1*$^{Cre}$, ns: not statistically significant.

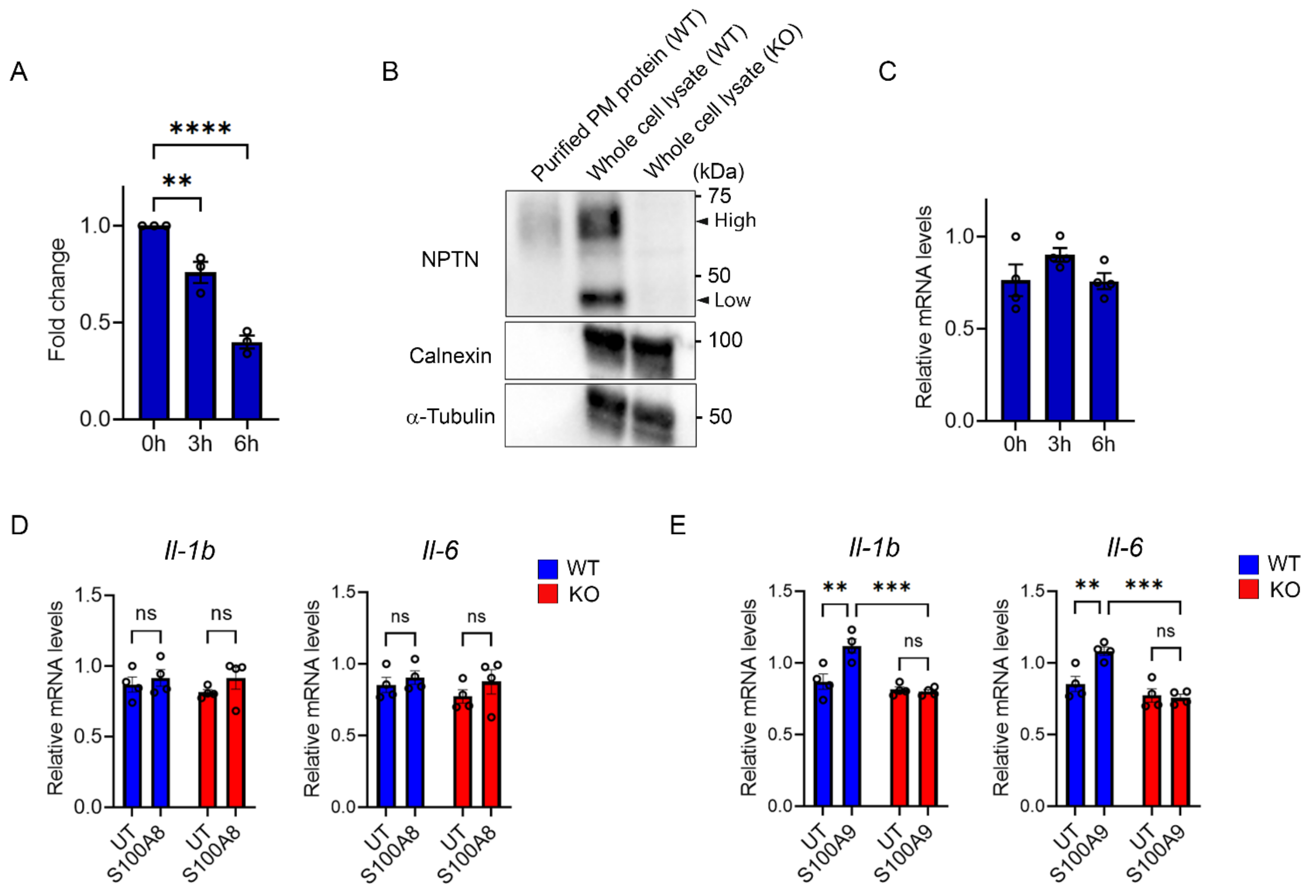


**Figure S6. Additional analyses of the NPTN roles in inflammation.**

(A) A quantification of NPTN protein levels at lower molecular weight in Figure 4J normalized to α-Tubulin (n=3, **p<0.01 and ****p<0.0001 by one-way ANOVA). (B) Representative blotting images of NPTN, ER marker (Calnexin), and α-Tubulin in the proteins purified from plasma membrane (PM) of *Nptn* WT, whole cell lysates extracted from *Nptn* WT or KO INS-1 832/13 cells. Note that NPTN signal at low molecular weight was not detected in purified PM proteins. (C) qPCR analyses of *Nptn* in INS-1 832/13 cells treated with cytokine mix (5 ng/ml IL-1β + 100 ng/ml IFNγ) for indicated times (n=4). (D-E) qPCR analyses of NF-κB target genes related to pro-inflammatory cytokines in *Nptn* WT or KO INS-1 832/13 cells treated with or without either 5 μg/ml S100A8 (D) or 5 μg/ml S100A9 (E) for 6 hours (n=4, **p<0.01 and ***p<0.001 by two-way ANOVA). ns: not statistically significant.

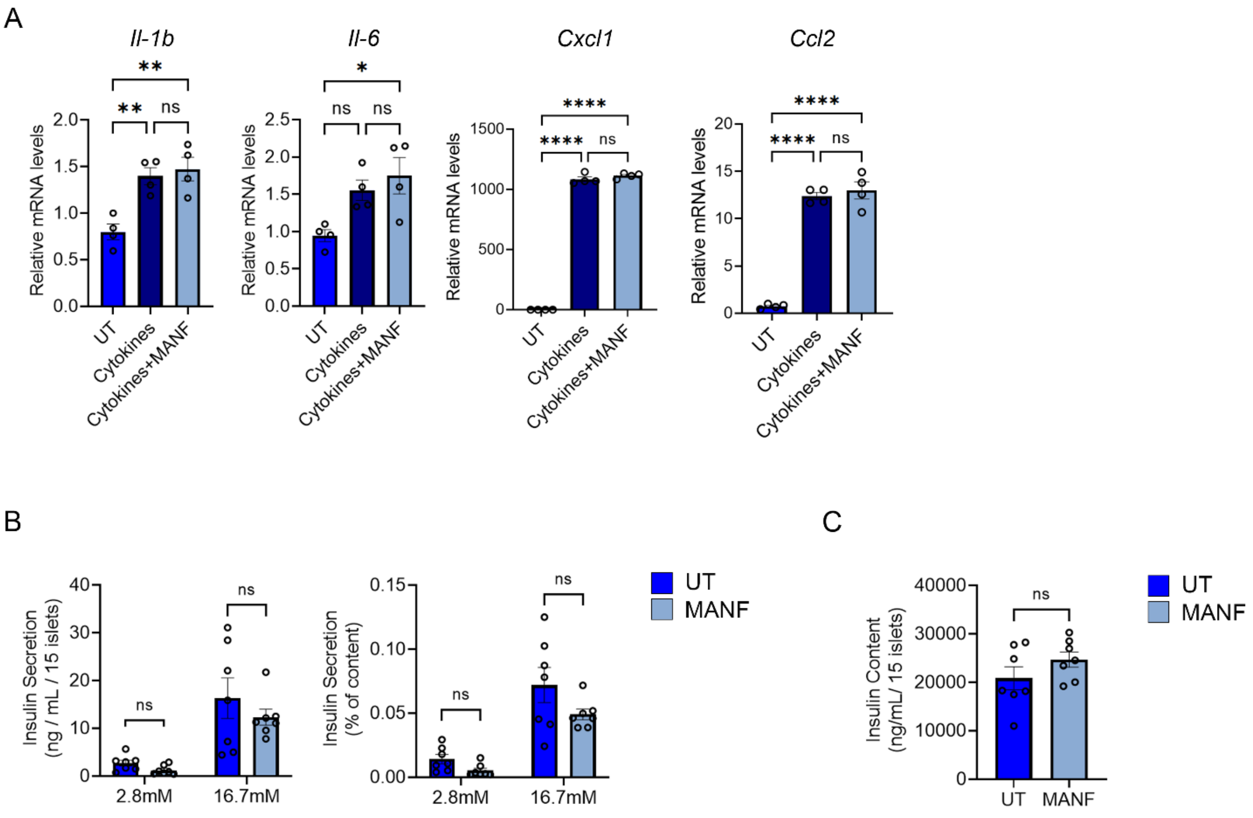


**Figure S7. Additional analyses of MANF effects on the NPTN-mediated inflammation and insulin secretion.**

(A) qPCR analysis of NF-κB target genes related to pro-inflammatory cytokines in *Nptn* KO INS-1 832/13 treated with or without cytokine mix (5 ng/ml IL-1β + 100 ng/ml IFNγ) and MANF (5 µg/ml) for 6 hours (n=4, *p<0.05, **p<0.01 and ****p<0.0001 by one-way ANOVA). (B) Static GSIS per islets (left) and normalized to insulin content (right) in primary islets form *Nptn*$^{f/f}$; *Ins1*$^{Cre}$ mice treated with or without MANF for 5 days (UT: n=7, MANF: n=7). (C) Insulin content in the islets used in (D). UT: untreated, ns: not statistically significant.

**Table S1. Antibody list used in this study.**

| Antibody | Company | Catalog # | Dilution |
|---|---|---|---|
| Insulin | CST | 3014 | 1:100 |
| Insulin | eBioscience | 53-9769-82 | 1:100 |
| Glucagon | Abcam | ab10988 | 1:100 |
| Neuroplastin | R&D | AF7818 | WB, 1:1000<br>IF, 1:100<br>IP, 1 μg |
| Ki67 | CST | 9129 | 1:100 |
| PMCA2 | Thermo | PA1-915 | 1:1000 |
| pDRP1[Ser637] | CST | 4867 | 1:1000 |
| DRP1 | CST | 8570 | 1:1000 |
| IκBα | CST | 4814 | 1:1000 |
| TRAF6 | Thermo | 38-0900 | 1:250 |
| Calnexin | Abcam | ab22595 | 1:1000 |
| IBA-1 | Novus | NB100-1028 | 1:50 |
| α-Tubulin | CST | 2125 | 1:1000 |

**Table S2. Primer list used in this study.**

| Primer | Species | Forward | Reverse |
|---|---|---|---|
| *Nptn* | Mouse | GAACGAACCAAGAATTGTCACCA | AGTGAGTTCTACCCCATTCCTT |
| *Nptn* | Rat | GCGCCAGAGAAACACAAATTAAG | GCATGCTTTAGACGGTCATTG |
| *Pcna* | Mouse | TACAGCTTACTCTGCGCTCC | TTGGACATGCTGGTGAGGTT |
| *Ccne2* | Mouse | ATGTCAAGACGCAGTAGCCG | CTGGGTTTCTTGCGGAGAGT |
| *Ccna2* | Mouse | GCCTTCACCATTCATGTGGAT | TTGCTCCGGGTAAAGAGACAG |
| *Cdk2* | Mouse | AGAAGATTGGAGAGGGCACG | ACACCTTCAGTCTCAGTGTCG |
| *Pola2* | Mouse | GAAACTGGCAGAGCTGTGTG | AGTAAGGCATGTCTTCCCCG |
| *Pdx1* | Mouse | CAGTGGGCAGGAGGTGCTTA | GGGCCGGGAGATGTATTTGTT |
| *Nkx6.1* | Mouse | GGCTGTGGGATGTTAGCTGT | TCATCTCGGCCATACTGTGC |
| *MafA* | Mouse | CAAGGAGGAGGTCATCCGAC | TCTCCAGAATGTGCCGCTG |
| *mtDNA* | Mouse | TTAAGACACCTTGCCTAGCCACAC | CGGTGGCTGGCACGAAATT |
| *nDNA* | Mouse | ATGACGATATCGCTGCGCTG | TCACTTACCTGGTGCCTAGGGC |
| *Il-1b* | Rat | CTCTGTGACTCGTGGGATGA | CGAGGCATTTTTGTTGTTCA |
| *Il-6* | Rat | TAGTCCTTCCTACCCCAACTTCC | TTGGTCCTTAGCCACTCCTTC |
| *Cxcl1* | Rat | CATTAATATTTAACGATGTGGATGCGTTTCA | GCCTACCATCTTTAAACTGCACAAT |
| *Ccl2* | Rat | CTTCTGGGCCTGTTGTTCAC | GCCAGTGAATGAGTAGCAGC |
| *18SrRNA* | Mouse/ Rat | AGGTTTGTGATGCCCTTAGATGTC | CACACGCTGAGCCAGTCAGT |
